# Physiologic Doses of TGF-β Improve the Composition of Engineered Articular Cartilage

**DOI:** 10.1101/2023.09.27.559554

**Authors:** Tianbai Wang, Sedat Dogru, Zhonghao Dai, Sung Yeon Kim, Nicholas A. Vickers, Michael B. Albro

## Abstract

For cartilage regeneration applications, transforming growth factor beta (TGF-β) is conventionally administered at highly supraphysiologic doses (10-10,000 ng/mL) in an attempt to cue cells to fabricate neocartilage that matches the composition, structure, and functional properties of native hyaline cartilage. While supraphysiologic doses enhance ECM biosynthesis, they are also associated with inducing detrimental tissue features, such as fibrocartilage matrix deposition, pathologic-like chondrocyte clustering, and tissue swelling. Here we investigate the hypothesis that moderated TGF-β doses (0.1-1 ng/mL), akin to those present during physiological cartilage development, can improve neocartilage composition. Variable doses of media-supplemented TGF-β were administered to a model system of reduced-size cylindrical constructs (Ø2-Ø3 mm), which mitigate the TGF-β spatial gradients observed in conventional-size constructs (Ø4-Ø6 mm), allowing for a novel assessment of the intrinsic effect of TGF-β doses on macroscale neocartilage properties and composition. The administration of physiologic TGF-β to reduced-size constructs yields neocartilage with native-matched sGAG content and mechanical properties while providing a more hyaline cartilage-like composition, marked by: 1) reduced fibrocartilage-associated type I collagen, 2) 77% reduction in the fraction of cells present in a clustered morphology, and 3) 45% reduction in the degree of tissue swelling. Physiologic TGF-β appears to achieve an important balance of promoting requisite ECM biosynthesis, while mitigating hyaline cartilage compositional deficits. These results can guide the development of novel physiologic TGF-β-delivering scaffolds to improve the regeneration clinical-sized neocartilage tissues.

## Introduction

Cartilage tissue engineering is a promising osteoarthritis (OA) treatment strategy for the surgical repair of clinical chondral defects [1]. Conventionally, regenerative platforms consist of the initial *in vitro* cultivation of cell-seeded scaffolds in an attempt to regenerate neocartilage that recapitulates the composition, structure, and functional mechanical properties of hyaline cartilage. This consists of: 1) an extracellular matrix (ECM) comprised of a dense array of sulfated glycosaminoglycans (sGAG) enmeshed in a type II collagen (Col-II) network [2], which gives rise to the tissue’s capacity to support physiologic mechanical loads, and 2) a sparse array of isolated chondrocytes, which actively synthesize and replenish the ECM constituents. To date, the field has been burdened by limited clinical outcomes, marked by short term survival of repair tissue after clinical implantation [3]. These outcomes are believed to result from neocartilage with inferior ECM composition and mechanical properties, thus supporting the need for the development of improved regenerative platforms.

Transforming growth factor beta (TGF-β) is a potent anabolic hormone that has emerged as one of the most widely utilized mediators in cartilage tissue engineering due to its ability to promote chondrogenesis and cartilage ECM biosynthesis [4, 5]. During periods of *in vitro* tissue cultivation, TGF-β is supplemented in culture medium to stimulate cell biosynthesis. Additionally, in recent years, an assortment of scaffold-based TGF-β delivery platforms have been developed (e.g., affinity domains, microspheres, liposomes) that offer continued delivery of TGF-β *in vivo* after clinical implantation [6]. In general, for these regenerative platforms, TGF-β is conventionally administered to cells at highly supraphysiologic doses, ranging between 10-10,000 ng/mL [4, 7-19] (Table 1). While these doses accelerate tissue growth, interestingly, they have long been associated with the induction of pathology in native connective tissues. TGF-β is significantly elevated in OA and contributes to the hallmark features of joint pathology, including hyperplasia, synovial fibrosis, and osteophyte formation [20]. Conventional supraphysiologic TGF-β doses in regenerative platforms can lead to similar tissue abnormalities, including: 1) the pronounced deposition of fibrocartilage-associated type I collagen (Col-I), in contrast to Col-II characteristic of hyaline cartilage [21], 2) the induction of cell hyperplasia, marked by formation of dense cell clusters [22], reminiscent of the abnormal chondrocyte morphology observed in OA [23], and 3) severe tissue swelling as a result of accelerated sGAG biosynthesis in the presence of a slowly regenerating collagen network [24].

**Table 1.** Administered doses of active TGF-β in tissue engineering platforms.

| Delivery platform | Delivery modality | Biomaterial scaffold | Loaded TGF-β dose | References |
| --- | --- | --- | --- | --- |
| Media supplementation | Culture media | N/A | 10 ng/mL | [4, 7, 8] |
|  |  |  | 30 ng/mL | [9, 10] |
|  |  |  | 100 ng/mL | [11] |
| Scaffold releasing | Microspheres | Alginate | 2 mg/mL | [12] |
|  |  | Gelatin | 3-6 mg/mL | [13] |
|  | Scaffold tethering | Chitosan and collagen | 10 mg/mL | [14] |
|  |  | Polyethylene glycol (PEG) | 250-2,500 ng/mL | [15] |
|  | Scaffold encapsulation | Self-assembling peptide | 100 ng/mL | [16] |
|  |  | Poly-D,L-lactic acid (PDLLA)-PEG and hyaluronic acid (HA) | 10 mg/mL | [17] |
|  | Affinity binding domains | Heparin and HA | 200 ng/mL | [18] |
|  |  | Alginate sulfate | 200 ng/mL | [19] |

In contrast to conventional dosing regimens during cartilage regeneration, TGF-β delivered to chondrocytes during native cartilage development occurs in a highly moderated fashion. Here, TGF-β is introduced to cells via its characteristic latent complex [25]—consisting of the active TGF-β peptide sequestered by a propeptide shell protein—which allows for slow, tightly controlled release (termed “activation”), via physiologic mechanical or enzymatic triggers. This mechanism provides developing cartilage with sufficient TGF-β exposure regimens needed to achieve requisite biosynthesis while mitigating pathogenesis associated with TGF-β excesses. For context, steady-state levels of activated TGF-β in the healthy synovial joint are generally low, in the range of 0.01 to 0.7 ng/mL [26-29] (Table 2), representing levels that are one to six orders of magnitude below those administered during conventional cartilage regeneration protocols. When considering this developmental paradigm, it may be reasonable to surmise that the administration of TGF-β at moderated, physiologic-range doses may give rise to improved cartilage regeneration outcomes.

**Table 2.** Measured levels of active TGF-β in healthy and diseased articular cartilage and synovial fluid. OA: osteoarthritis; RA: rheumatoid arthritis.

| Condition | Tissue type | Species | Active TGF-β level | Measurement technique | References |
| --- | --- | --- | --- | --- | --- |
| Healthy | Articular cartilage | Bovine | 0.7 ng/mL | ELISA | [26] |
|  | Articular cartilage | Bovine | 0.15 ng/mL | Bioassay | [27] |
|  | Synovial fluid | Bovine | 0.01-0.05 ng/mL | ELISA | [28] |
|  | Synovial fluid | Human | 0.03 ng/mL | ELISA | [28] |
| Diseased | Synovial fluid (OA) | Human | 3.8 ng/mL | Radioreceptor assay | [29] |
|  | Synovial fluid (RA) | Human | 10.1 ng/mL | Radioreceptor assay | [29] |

Interestingly, studies of tissue growth in response to physiologic TGF-β doses are limited. Byers et al. [4] monitored *in vitro* growth of Ø5×2.3 mm engineered constructs in response to a range of media-supplemented TGF-β doses, but found that tissue mechanical properties at a near-physiologic dose (1 ng/mL) were significantly lower than those achieved at a supraphysiologic dose (10 ng/mL), thus suggesting that physiologic TGF-β dosing may in fact be insufficient to provide requisite rates of ECM biosynthesis and supporting the use of supraphysiologic doses as a baseline standard for the field. However, recently developed insights suggest that this interpretation may be incomplete, as it does not account for the previously unanticipated complexity of spatial distribution profiles of administered TGF-β in constructs. To this end, our recent study has demonstrated that TGF-β exhibits pronounced steady state gradients in conventional-size tissue constructs due to pronounced rates of reversible binding to the construct scaffold and consumption by embedded cells [22]. As a result, one is unable to ascertain whether the low growth profiles observed for physiologic TGF-β doses result from their lower intrinsic biosynthetic effect or rather an artifact from their reduced penetration into conventional-size tissues.

Based on these new insights, we aim to revisit the fundamental question of whether the administration of physiologic doses of TGF-β can improve neocartilage regeneration. Specifically, we hypothesize that, in the absence of spatial distribution artifacts, physiologic doses of TGF-β can yield neocartilage with a more hyaline-cartilage-like composition and structure compared to supraphysiologic doses. To explore this hypothesis, we compare the effect of media-supplemented physiologic and conventional supraphysiologic TGF-β doses on the growth of neocartilage using a reduced-size construct (Ø2-Ø3 mm) model system, which exhibits mitigated TGF-β spatial gradients, allowing for a more faithful evaluation of the intrinsic effect of TGF-β doses on promoting cartilage regeneration. Using this model, we explore the capability of physiologic doses to yield macroscale neocartilage with native-matched mechanical properties while mitigating the hyaline cartilage quality deficits associated with supraphysiologic TGF-β doses, notably tissue swelling, elevated fibrocartilage-associated Col-I deposition, and hyperplasia-induced cell clustering. Potential regeneration benefits derived from physiologic TGF-β dose stimulation can guide the development and optimization of emerging TGF-β scaffold delivery platforms, allowing for the regeneration of improved quality larger sized constructs that are ultimately needed to achieve tissue repair in the clinic.

## Results

### Reducing construct size mitigates TGF-β concentration gradients

The ability of reduced-size constructs to mitigate spatial gradients of media-supplemented TGF-β and resulting heterogeneities of mechanical properties relative to conventional-size constructs was initially assessed. Conventional-size (Ø6×3 mm) bovine-chondrocyte-seeded agarose constructs were exposed to TGF-β 3 supplemented in medium at a physiologic (0.3 ng/mL) or a conventional supraphysiologic (10 ng/mL) dose for 5 days. Subsequently, constructs were axially sub-punched and sectioned, allowing for ELISA-based measures of the 1D gradient of TGF-β3 through the construct depth [22, 26]. Pronounced TGF-β concentration gradients were observed for both doses; TGF-β levels decreased to near zero at 0.6 and 1.1 mm below the tissue surface for 0.3 and 10 ng/mL, respectively (Fig. 1a). Subsequently, a finite element (FE) reaction diffusion model was implemented to describe the spatial distribution of TGF-β in conventional-size (Ø5×2 mm) and reduced-size (Ø2×2 mm) constructs, using previously measured parameters (Supplementary Table 1) [22]. Consistent with experimental measures, models predicted strong steady state TGF-β gradients in conventional-size constructs. However, in reduced-size constructs, gradients were significantly mitigated (Fig. 1b). The degree of construct ECM heterogeneity after 56 days of cultivation was further assessed. For conventional-size constructs, enhancements in sGAG and mechanical properties in response to 0.3 ng/mL and 10 ng/mL TGF-β doses occurred predominantly at the construct periphery, as demonstrated by biochemical assay measures of the 1D depth-dependent distribution of sGAG (Fig. 1c) and digital image correlation (DIC) analysis of the strain distribution through the construct depth (Fig. 1d-e). In contrast, reduced-size constructs exhibited more homogeneous mechanical properties as demonstrated by increased uniformity in DIC strain profiles (Fig. 1d-e). These demonstrations support the use of reduced-size macroscale constructs to examine the intrinsic effect of varying TGF-β doses on construct growth outcomes.

**Fig. 1.**
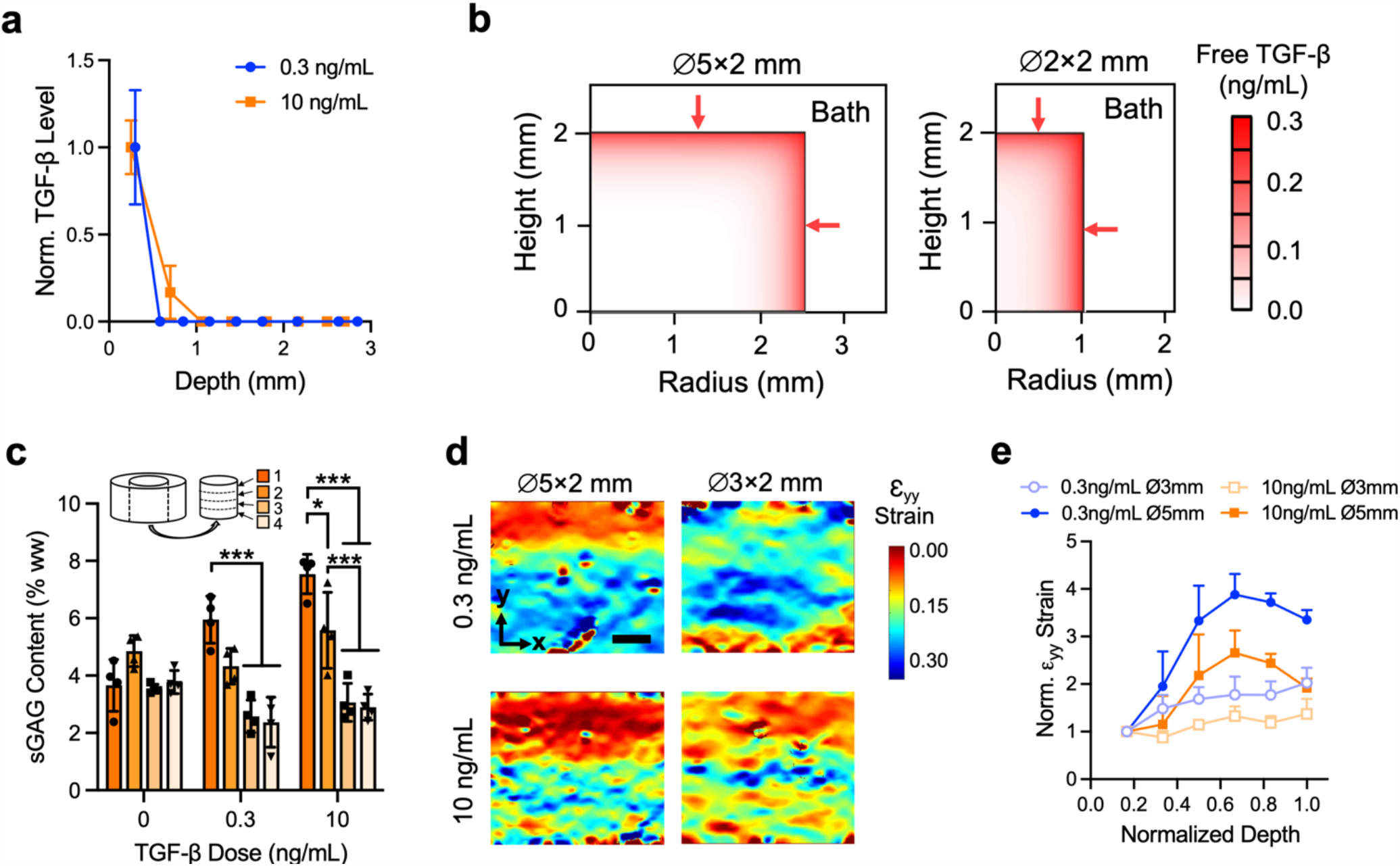
Mitigation of TGF-β concentration gradients and heterogeneities in reduced-size macroscale constructs. (**a**) Spatial distribution of media supplemented TGF-β through the depth of Ø6×3 mm tissue constructs after 5-day exposure of physiologic (0.3 ng/mL) or supraphysiologic (10 ng/mL) TGF-β dose. Error bars represent mean ± s.d. (**b**) FE model of distribution of media-supplemented TGF-β (0.3 ng/mL) at 72 hours for conventional-size (Ø5×2 mm) and reduced-size (Ø2×2 mm) construct (half cross-section depicted). Arrows represent the direction of TGF-β uptake. (**c**) Assay measured depth-dependent distribution of sGAG in Ø6×2 mm constructs at day 56 after exposure to 0, 0.3, or 10 ng/mL TGF-β. Error bars represent mean ± s.d. *p<0.05, ***p<0.001. (**d**) Representative DIC measured compressive strain (ε_yy_) distribution in response to 10% platen-to-platen strain for conventional-size (Ø5×2 mm) and reduced-size (Ø3×2 mm) constructs at day 56 after exposure to 0.3 or 10 ng/mL TGF-β. Scale bar: 500 mm. (**e**) Variation of normalized strain (ε_yy_) with tissue depth for Ø5×2 mm and Ø3×2 mm constructs exposed to 0.3 or 10 ng/mL TGF-β. Error bars represent mean + s.d.

### Physiologic TGF-β doses yield neocartilage with native-matched functional mechanical properties

We examined the effect of a range of media-supplemented TGF-β doses (TGF-β-free: 0 ng/mL; physiologic range doses: 0.1, 0.3, 1 ng/mL; supraphysiologic range doses: 3, 10 ng/mL) on the mechanical properties of conventional-size constructs (Ø4-Ø6×2 mm), which exhibit pronounced TGF-β gradients, and reduced-size constructs (Ø2-Ø3×2 mm), which exhibit mitigated TGF-β gradients. Of interest, is the extent to which TGF-β exposure increases Young’s modulus toward native-cartilage values. After 56 days of culture, constructs were analyzed for their compressive Young’s modulus (E_Y_). Generally, construct E_Y_ increased with TGF-β dose and decreased with construct size (Fig. 2). For conventional-size constructs, physiologic range TGF-β doses induced only modest E_Y_ enhancements; E_Y_ failed to reach native levels for all doses and no significant increase above TGF-β-free levels was observed for 0.1 ng/mL and 0.3 ng/mL doses (p=1.0; Fig. 2a). In contrast, for reducedsize Ø2 mm constructs, physiologic doses (0.3 ng/mL and 1 ng/mL) yielded constructs with native-matched E_Y_ (Fig. 2b), indicating that, when sufficient penetration into the tissue is achieved, physiologic doses are capable of recapitulating functional mechanical properties characteristic of healthy cartilage.

**Fig. 2.**
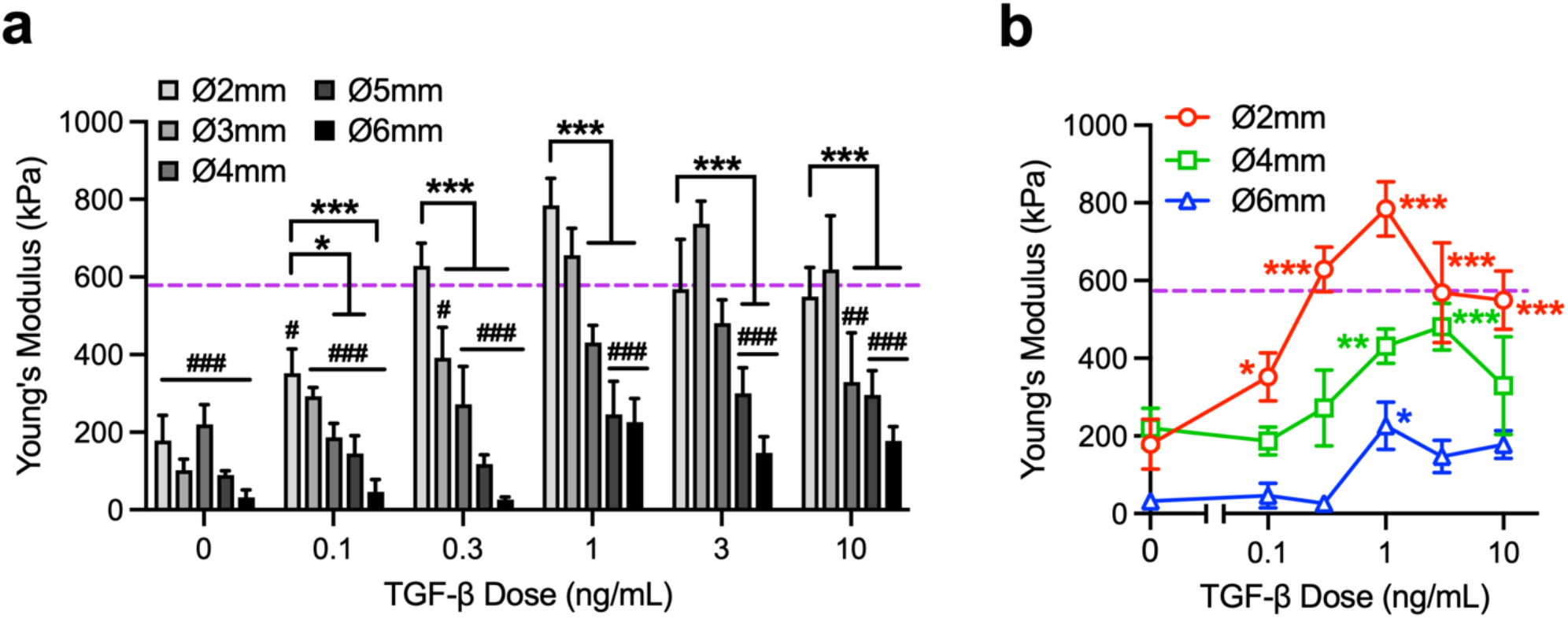
Neocartilage mechanical properties in response to varying TGF-β doses. (**a**) Compressive Young’s modulus (E_Y_) of reduced-size (Ø2– Ø3×2 mm) and conventional-size (Ø4– Ø6×2 mm) constructs at day 56 in response to 0 ng/mL, physiologic (0.1-1 ng/mL), or supraphysiologic (3-10 ng/mL) TGF-β doses. Dashed line represents average E_Y_ of native bovine cartilage (590±150 kPa). Error bars represent mean + s.d. *p<0.05, ***p<0.001. ^#^p<0.05, ^##^p<0.005, ^###^p<0.001: significantly below the native E_Y_ level. (**b**) E_Y_ of Ø2×2 mm, Ø4×2 mm, and Ø6×2 mm constructs at day 56 in response to all TGF-β doses. Dashed line represents average E_Y_ of native bovine cartilage. Error bars represent mean ± s.d. *p<0.05, **p<0.005, ***p<0.001: significant increase above the 0 ng/mL level.

### Physiologic TGF-β doses yield neocartilage with mitigated tissue swelling

Healthy articular cartilage exists in an equilibrium state wherein the sGAG induces a Donnan osmotic swelling pressure that is balanced by the tensile restraining properties of the collagen matrix [30, 31]. Developing neocartilage rapidly accumulates sGAG but possesses a slowly developing collagen matrix causing the tissue to progressively swell during development [24]. Supraphysiologic TGF-β doses can further exacerbate swelling by accelerating sGAG biosynthesis. Here, we measured the sGAG content, collagen content, and degree of swelling of constructs in response to physiologic and supraphysiologic TGF-β doses. The sGAG and collagen contents generally decrease with construct size (Fig. 3a and 3b), in a manner akin to E_Y_. For Ø2 mm constructs, sGAG content increased with TGF-β dose, reaching native levels at 0.3 ng/mL (p=0.97; Fig. 3c). Collagen content exhibited a biphasic response, increasing with TGF-β dose in the physiologic range and peaking at 1 ng/mL, but decreasing beyond in the supraphysiologic range (Fig. 3d). For the conventional supraphysiologic 10 ng/mL dose, the sGAG-to-collagen content ratio reached a high value of 10.2±1.7, in far excess of the ratio found in native cartilage (0.9±0.2; Fig. 3e). For physiologic doses, the sGAG-to-collagen content ratio was significantly lowered, reaching a value of 6.2±0.6 at 0.3 ng/mL (p<0.001). Construct swelling was quantified with a swelling ratio (SR), relating the construct final mass to its initial mass. For 10 ng/mL, significant swelling was observed (SR=2.4±0.3) (Fig. 3f). For physiologic doses, the SR was significantly lowered (p<0.001), reaching a value of only SR=1.3±0.1 for 0.3 ng/mL. The sGAG-to-collagen content ratio predicted 57% of the variation in SR (Fig. 3g). Overall, these results demonstrate that physiologic TGF-β doses can yield neocartilage with a more native-like sGAG-to-collagen content ratio, leading to a pronounced mitigation of tissue swelling during development.

**Fig. 3.**
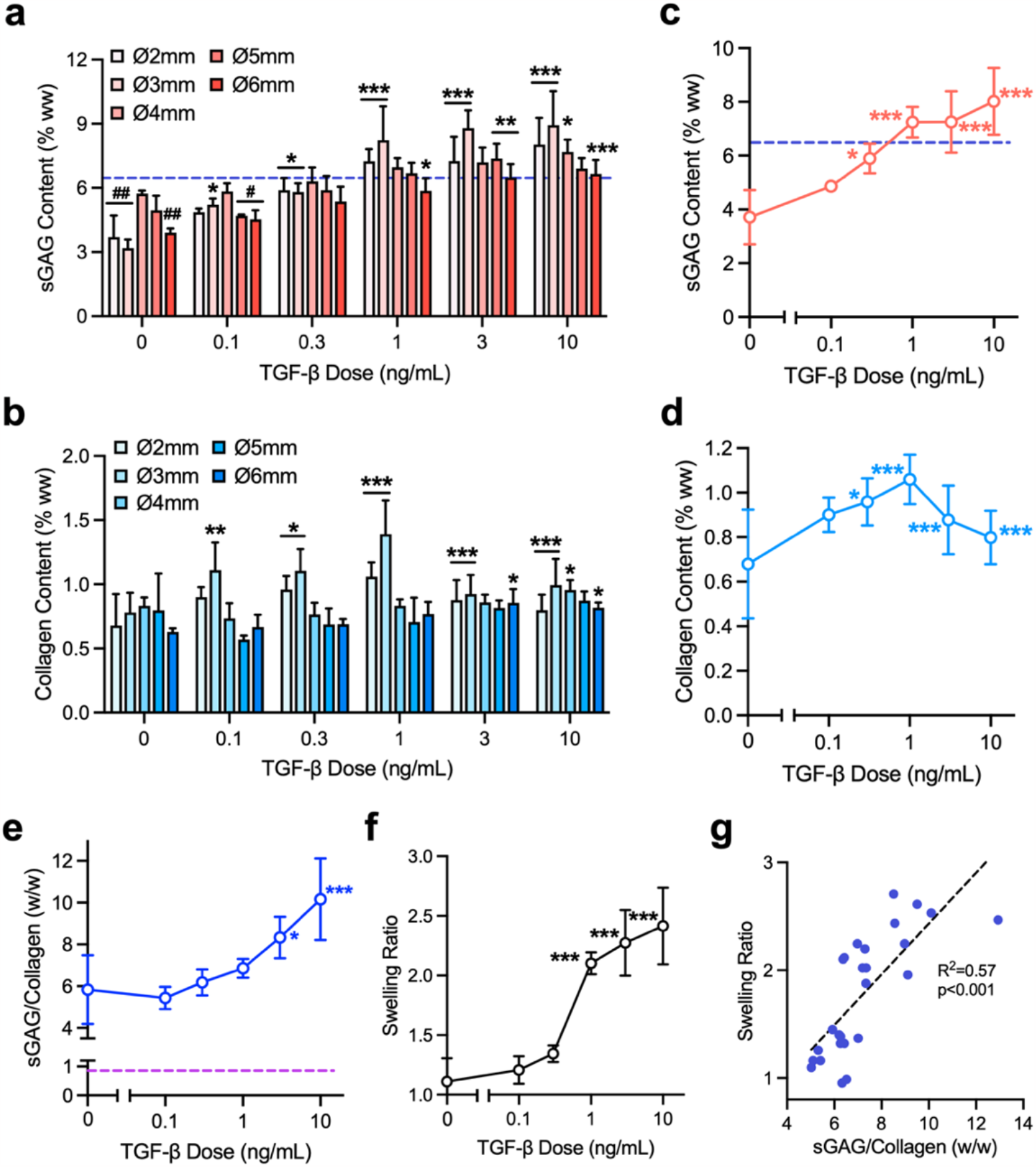
Neocartilage composition in response to varying doses of TGF-β. (**a**) sGAG and (**b**) total collagen contents of reduced-size (Ø2– Ø3×2 mm) and conventional-size (Ø4– Ø6×2 mm) constructs at day 56 in response to 0 ng/mL, physiologic (0.1-1 ng/mL), or supraphysiologic (3-10ng/mL) TGF-β doses. Dashed line represents average sGAG content of native bovine cartilage (6.5±1.6%ww). Error bars represent mean + s.d. (**c**) sGAG and (**d**) total collagen contents of reduced-size (Ø2×2 mm) constructs at day 56 in response to all TGF-β doses. Dashed line represents average sGAG content of native bovine cartilage. (**e**) Ratio of sGAG to total collagen content and (**f**) swelling ratio of Ø2×2 mm constructs in response to all TGF-β doses. Dashed line represents average sGAG to collagen ratio of native bovine cartilage (0.87±0.15). Error bars represent mean ± s.d. (**g**) Bivariate linear regression of sGAG-to-collagen content ratio versus swelling ratio for Ø2×2 mm constructs. *p<0.05, **p<0.005, ***p<0.001: significantly above the 0 ng/mL levels. ^#^p<0.05, ^##^p<0.005: significantly below the native sGAG content level.

### TGF-β doses yield biphasic neocartilage growth response

We next examined construct growth in response to a wider range of TGF-β doses with the inclusion of highly supraphysiologic doses that are conventionally administered for TGF-β scaffold delivery platforms (≥100 ng/mL) [6]. Here, reduced-size (Ø3 mm) constructs were exposed to media-supplemented TGF-β at 0, 0.1, 0.3, 1, 3, 10, 30, 100, or 300 ng/mL and analyzed for E_Y_ and sGAG after 14, 28, 42, or 56 days of culture, and DNA content at day 56. For all time points, construct E_Y_ exhibited a biphasic response, increasing with TGF-β dose until 3-10 ng/mL and then decreasing at doses beyond (Fig. 4a). Similar biphasic responses were observed for sGAG content (Fig. 4b) and DNA-based cell density (Fig. 4c). Overall, these results highlight the potential for highly supraphysiologic TGF-β doses to suppress the deposition of cartilage ECM, leading to neocartilage with inferior functional mechanical properties.

**Fig. 4.**
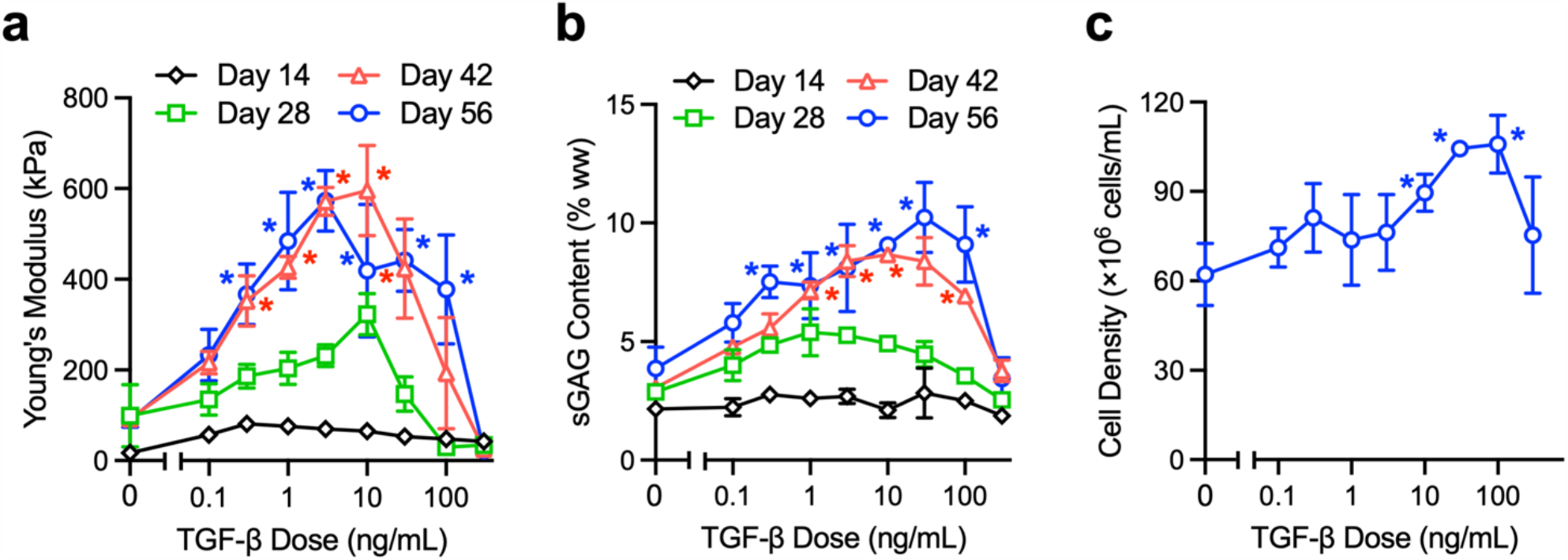
Biphasic effect of highly supraphysiologic TGF-β dose range on neocartilage mechanical properties and composition. (**a**) E_Y_ and (**b**) sGAG content of reduced-size constructs (Ø3×2 mm) at day 14, 28, 42, and 56 after exposure to a range of TGF-β doses. (**c**) Cell density of Ø3×2 mm constructs at day 56 after exposure to a range of TGF-β doses. All error bars represent mean ± s.d. *p<0.05: significant increase above corresponding 0 ng/mL levels.

### Physiologic TGF-β doses yield neocartilage with mitigated Col-I deposition

A significant challenge in articular cartilage regeneration is promoting the deposition of hyaline cartilage-associated Col-II while mitigating the deposition of fibrocartilage-associated Col-I. Here, we assessed whether physiologic TGF-β doses can reduce levels of Col-I in constructs. Immunostaining for Col-I and Col-II was performed on reduced-size (Ø3 mm) constructs after 56 days of culture. For Col-II, similar staining intensities were observed for all TGF-β doses (Fig. 5a, bottom row). For Col-I, staining was found in both intracellular and extracellular regions of the constructs exposed to supraphysiologic TGF-β doses (10, 100, 300 ng/mL), but was greatly reduced for the physiologic TGF-β dose at 0.3 ng/mL (Fig. 5a, top row). Further, the quantified results demonstrated significantly lower Col-I positive cells in constructs exposed to physiologic TGF-β doses than supraphysiologic doses (p<0.001, Fig. 5b), supporting the ability of physiologic TGF-β dosing to promote a more hyaline-cartilage-like tissue composition.

**Fig. 5.**
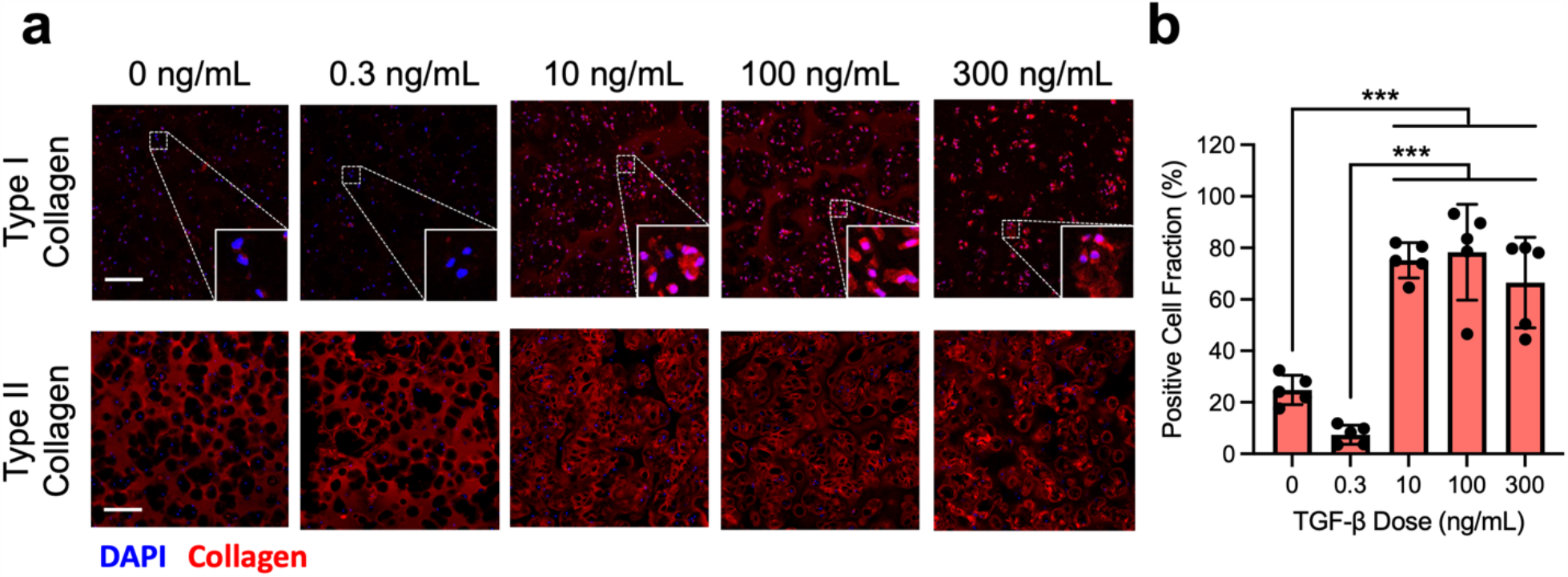
Deposition of collagen subtypes in response to varying doses of TGF-β. (**a**) Immunofluorescence overlay of type I/II collagen (red) and chondrocyte nuclei (blue) for reduced-size constructs (Ø3×2 mm) at day 56 after exposure to 0 ng/mL, physiologic (0.3 ng/mL), or supraphysiologic (10, 100, 300 ng/mL) TGF-β doses. Scale bar: 100 mm. (**b**) Fraction of cells positively stained for type I collagen. Error bars represent mean ± s.d. ***p<0.001.

### Physiologic TGF-β doses yield neocartilage with mitigated chondrocyte clustering

A hallmark feature of OA is the emergence of chondrocytes in dense clusters in contrast to their isolated morphology that is characteristic of healthy hyaline cartilage [23]. We next examined the dose-dependent effect of TGF-β on the morphology of chondrocytes in neocartilage. Reduced-size (Ø3 mm) constructs were exposed to varying TGF-β doses (0-10 ng/mL), stained with calcein-AM, and imaged via confocal microscopy after 30 and 56 days of culture (Fig. 6a). An image processing algorithm was developed to identify cellular regions with an isolated or clustered morphology (Supplementary Fig. 1-5), quantified as a cell cluster area fraction (CAF; ratio between area of cells in a clustered morphology and total cell area) (Fig. 6b). At day 0, only 1% of construct cells were present in a clustered morphology (CAF=0.01±0.01) (Fig. S4), akin to cluster levels in healthy native bovine cartilage (CAF=0.01±0.02). At day 30, cell clustering increased with TGF-β dose (p<0.001). For the conventional supraphysiologic 10 ng/mL dose, 58% of cells were present in clusters (CAF=0.58±0.02). Physiologic TGF-β doses mitigated cell clustering. At 0.1 ng/mL and 0.3 ng/mL doses, cell clusters levels (10% and 13%, respectively) were not significantly elevated above TGF-β-free conditions (p>0.85). Further, for 10 ng/mL, cell clustering increased further by day 56 (p<0.001), while cluster levels remained stable for 0.3 ng/mL (p=0.41). The 3D reconstructed confocal images further illustrate the morphology of cells in constructs for different TGF-β doses (Fig. 6c). Overall, these results highlight the ability of physiologic TGF-β doses to improve the morphology of chondrocytes during cartilage regeneration.

**Fig. 6.**
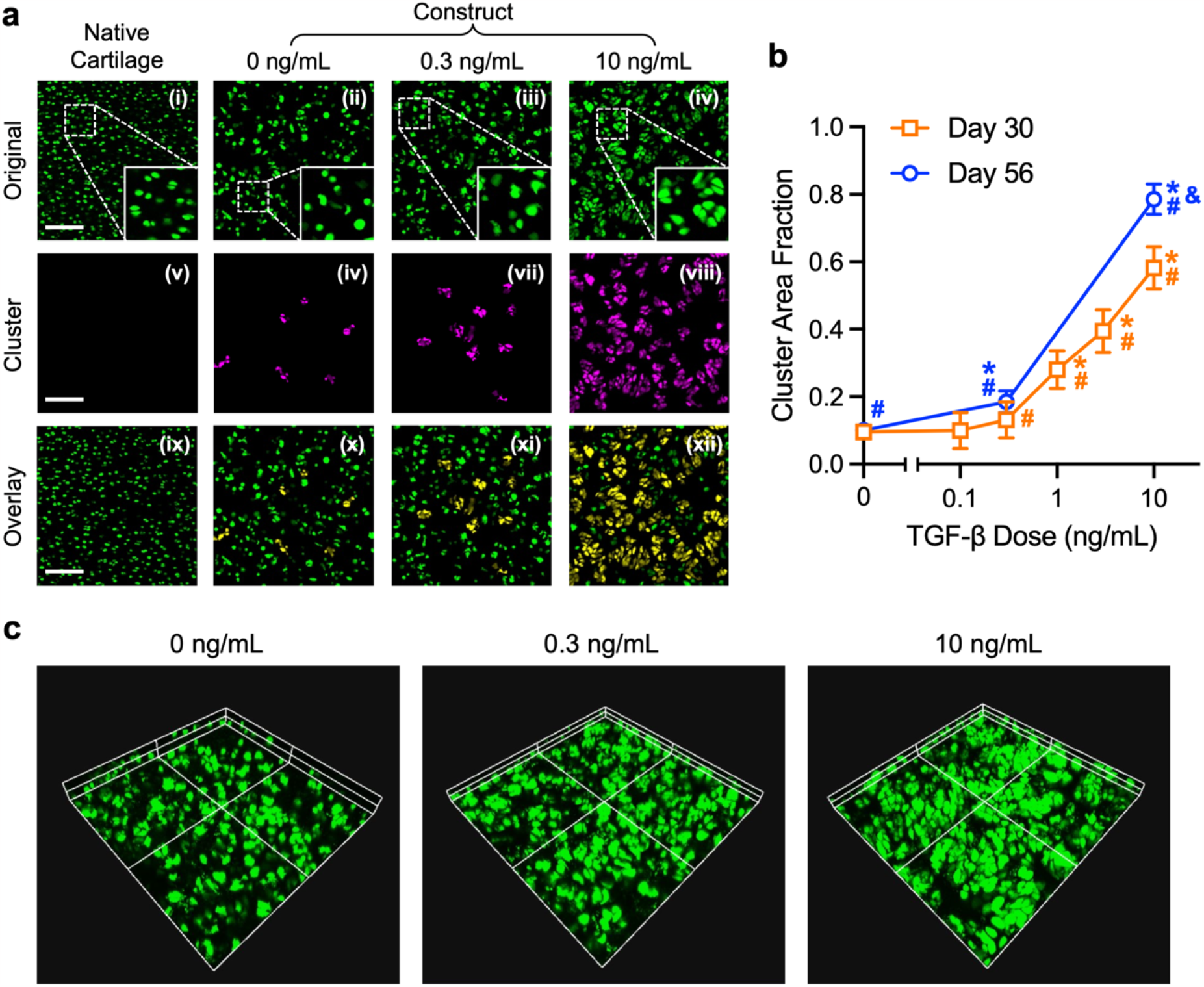
Chondrocyte morphology in response to varying TGF-β doses. (**a**) Representative images of cell morphology analysis for native bovine cartilage, and reduced-size constructs (Ø3×2 mm) at day 56 after exposure to 0 ng/mL, physiologic (0.3 ng/mL), or supraphysiologic (10 ng/mL) TGF-β doses. (i-iv) Calcein-AM stained chondrocytes (green), (v-viii) identification of clustered chondrocytes (magenta) through image analysis, and (ix-xii) overlay of clustered chondrocytes (yellow) and isolated chondrocytes (green). Scale bar: 150 mm. (**b**) Cell cluster area fraction (CAF) of constructs at days 30 and 56 after exposure to a range of TGF-β doses. CAF of native cartilage and day 0 constructs is at 0.01±0.02 and 0.01±0.01, respectively. Error bars represent mean ± s.d. * ^#^ & p<0.05: significantly above the corresponding 0 ng/mL TGF-β, native, and day 30 levels, respectively. (**c**) 3D views of cells in Ø3×2 mm constructs exposed to 0, 0.3, or 10 ng/mL TGF-β after 56 days of culture.

### Physiologic TGF-β doses yield neocartilage with more homogenous cell strains

A functional consequence of TGF-β induced cell clusters may be alterations to the cellular mechanical environment created by physiologic mechanical loading after implantation. We performed a computational analysis to explore the influence of cell clustering on cell and tissue deformation in response to physiologic loading. Minimum principal stress and strain for ECM and cellular regions were predicted in response to 20 kPa axial compression (to maintain infinitesimal cell strains) via an FE model using experimentally derived cell morphology profiles from aforementioned confocal images (Fig. 6a). Models demonstrated that the clustered cellular regions derived from supraphysiologic 10 ng/mL TGF-β exposure exhibited an elevated and more disperse distribution of minimum principal compressive stress (σ=246±80 Pa) and strain (ε=0.33±0.11) relative to those of the more isolated cellular regions derived from physiologic 0.3 ng/mL TGF-β exposure (σ=177±50 Pa, ε=0.21±0.06) and in native cartilage (σ=203±46 Pa, ε=0.21±0.05) (Fig. 7a-b). Stress and strain in ECM regions exhibited similar trends (Fig. 7c). Overall, this analysis suggests that cells in neocartilage produced by supraphysiologic TGF-β will experience aberrant and heterogeneous strain profiles while cells in neocartilage produced by physiologic TGF-β will experience strains more akin to chondrocytes in healthy cartilage.

**Fig. 7.**
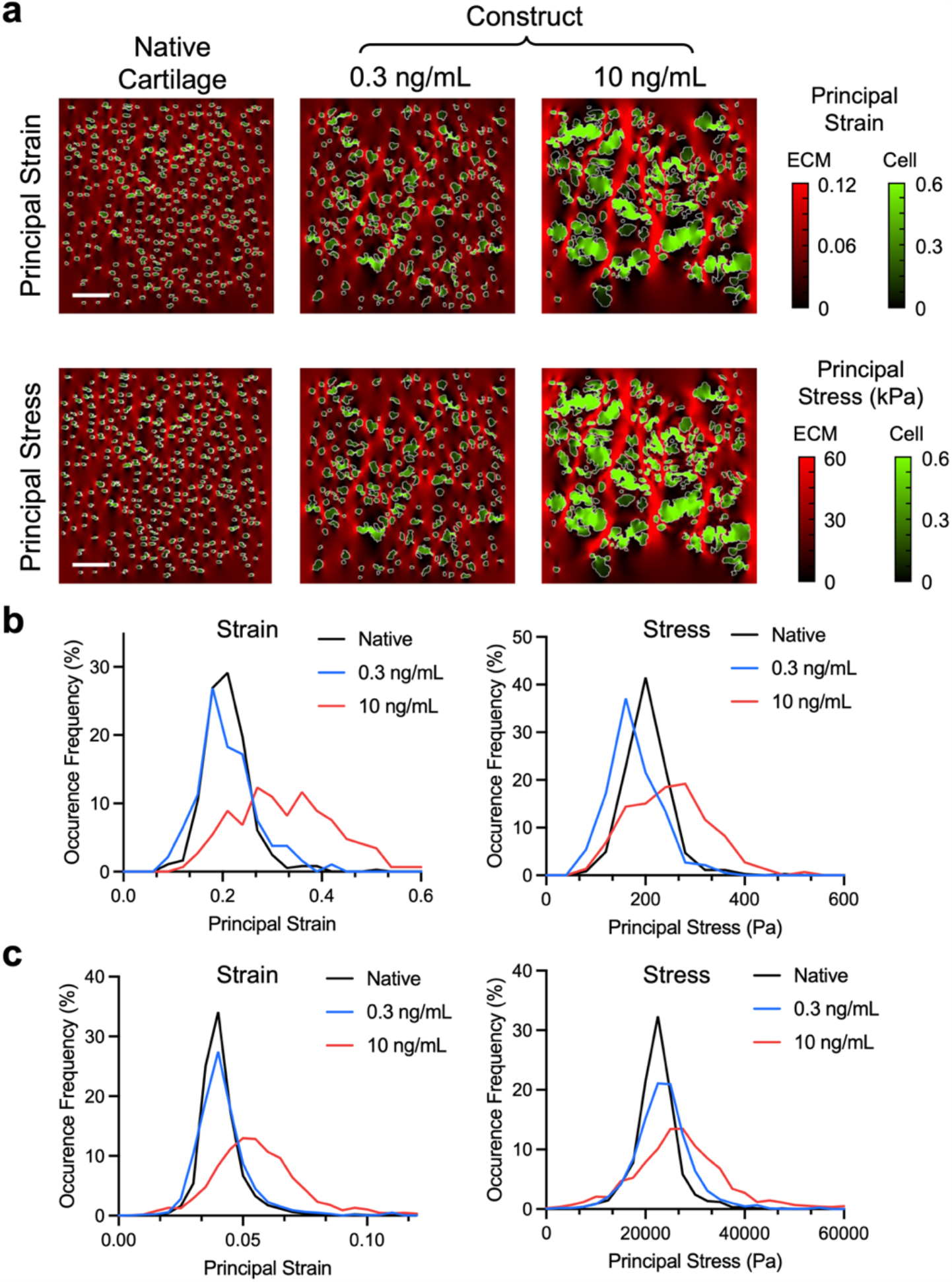
Predicted minimum principal strain and stress in cartilage tissue under physiologic loading. FE model predicted representative color maps (**a**) and distributions of minimum principal strain and minimum principal stress of cellular (**b**) and ECM (**c**) regions for day 56 tissue constructs exposed to physiologic (0.3 ng/mL) or supraphysiologic (10 ng/mL) TGF-β doses in response to 20 kPa compressive stress. In the color maps, green and red depicts cell and ECM regions, respectively. Scale bar: 100 mm. Results of native cartilage provided for reference.

## Discussion

The results of this study advance a new perspective for the field of cartilage tissue engineering, demonstrating that the administration of TGF-β at moderated, physiologic doses can improve neocartilage regeneration. A novel examination of the dose-dependent effect of TGF-β on neocartilage growth was facilitated using a reduced-size, macroscale tissue model system, whereby the effect of prescribed doses of TGF-β on neocartilage growth could be determined in the absence of confounding effects of spatial TGF-β concentration gradients. Consistent with prior work [22], characterizations demonstrate that TGF-β delivered via culture media supplementation exhibits pronounced steady-state gradients in constructs as a result of a high rate of consumption by embedded chondrocytes. For constructs of the size conventionally used in tissue engineering studies, TGF-β gradients can be highly pronounced—free TGF-β levels decay by ∼90% within 500 mm from the media-exposed surface, leading to heterogeneities in ECM deposition and tissue mechanical properties (Fig. 1). In contrast, for reduced-size constructs, media-supplemented TGF-β gradients are significantly mitigated, leading to a more uniform stimulation of embedded chondrocytes, as evidenced by improved homogeneity of construct properties (Fig. 1d-e). Overall, the reduced-size construct model system allows us to perform a foundational assessment of the intrinsic effect of TGF-β doses on the composition and mechanical properties of engineered neocartilage. Here, we observe that physiologic range TGF-β doses (0.3 ng/mL) can achieve an important balance of promoting ECM biosynthesis rates needed to yield neocartilage with native-matched mechanical properties, while mitigating the induction of commonly observed compositional abnormalities of regenerating cartilage, including tissue swelling, Col-I deposition, and chondrocyte clustering.

The recapitulation of a dense, functional collagen matrix for cartilage regeneration has remained a long-standing challenge. Generally, cell sources adopted for tissue engineering (e.g., chondrocytes, mesenchymal stem cells) synthesize sGAG at a significantly higher rate than collagen, an imbalance that is exacerbated by stimulation with supraphysiologic TGF-β doses. While the sGAG-rich ECM composition allows for a recapitulation of native-matched compressive properties, when combined with the slow-to-develop collagen matrix, a highly pronounced tissue swelling is observed. Here we highlight the severity of this imbalance in constructs exposed to supraphysiologic TGF-β doses, which exhibit a GAG-to-collagen mass ratio that is ∼10-fold higher than that observed in healthy native cartilage, causing the tissue to dramatically swell by 250% of its initial volume (Fig. 3e-f). The alternative administration of physiologic TGF-β doses yields neocartilage that still achieves native-matched compressive E_Y_, but while achieving a reduced, more native-like sGAG-to-collagen ratio, leading to mitigated swelling. It is important to note that the sGAG-to-collagen ratio remains above that present in functional native cartilage. In the future, physiologic TGF-β administration can be potentially combined with other collagen promoting strategies to yield further improvements [32-35]. Based on theoretical insights, the improved sGAG-to-collagen ratio may further improve the tensile properties, dynamic modulus, and lubricity of neocartilage [36], which will be examined in future work.

In addition to absolute levels of collagen content, the levels of specific collagen subtypes deposited during regeneration is also critical for long-term tissue survival. The emergence of fibrocartilage, marked by an elevated composition of Col-I, is a widely observed detrimental outcome for standard-of-care cartilage regeneration treatments (e.g., autologous chondrocyte implantation and microfracture [37]) that results in tissues comprised of a collagen network with inferior mechanical properties, accelerated rates of enzymatic degradation, and a reduced capability to retain intrafibrillar water [36]. Here, we observe that supraphysiologic doses of TGF-β (≥10 ng/mL) give rise to an elevated deposition of Col-I in constructs (Fig. 5), consistent with prior demonstrations of colocalization of Col-I in construct peripheral regions where supraphysiologic TGF-β is concentrated [21]. Alternatively, physiologic TGF-β doses mitigate Col-I deposition, giving rise to a more hyaline-cartilage-like tissue comprised predominantly of Col-II.

The emergence of dense chondrocyte clusters in response to supraphysiologic TGF-β is a striking outcome, particularly when considering their similarity to the morphology of pathologic chondrocyte clusters classically observed in OA cartilage [23]. In constructs, cell clusters likely result from TGF-β mediated acceleration of cell proliferation while in the confines of an agarose scaffold that restricts cell expansion and migration. This response is consistent with the well-documented effect of TGF-β excesses in inducing cell hyperplasia [38, 39]. Alternatively, physiologic TGF-β doses mitigate cell cluster formation, leading to an isolated cell morphology (Fig. 6), akin to that of chondrocytes in healthy articular cartilage. While the detrimental impact of cell clusters may not be immediately evident during short-term *in vitro* growth phases, it is reasonable to anticipate that they may compromise long-term tissue health after clinical implantation. For one, the clustered morphology may be marker of adverse cell phenotype changes [23, 40]. Further, the cell morphology changes are likely to impact the deformation profiles experienced by cells in response to *in situ* physiologic loading. FE analysis provides an estimate of this deformation (Fig. 7), illustrating that cell clusters experience strain profiles that are elevated and far more heterogeneous than those experienced by isolated cells. Given the high sensitivity of chondrocytes to strain magnitude [41], it is reasonable to surmise that clustered cells derived from supraphysiologic TGF-β may lose their ability to respond in unison to physiologic mechanical stimuli. In contrast, the more isolated cells derived from physiologic TGF-β may maintain their ability to provide a harmonious response to loading, thus better recapitulating the mechanobiological behavior needed to maintain long-term tissue health. In the future, the quantitative accuracy of model predictions can be improved by implementing models with 3D cell morphology measures and the adoption of advanced constitutive relations. Model predictions can further be validated by *in situ* experimental measures of cell strains in response to loading.

The beneficial effects of physiologic TGF-β doses for neocartilage development are consistent with a myriad of evolutionary regulatory mechanisms that exist to prevent excessive TGF-β activity in native tissues. Most notably, TGF-β is presented to cells in its latent complex, whereby activation is elicited by mechanical or enzymatic environmental triggers, thus ensuring that TGF-β activity is commensurate with biological need. Further, once activated, TGF-β is rapidly cleared from the extracellular space, attributed to the scavenging proteins (e.g., a2-macroglobulin [42]), giving rise to a half-life on the order of several minutes [43]. A breakdown of the regulation of TGF-β latency often serves as a precursor to pathology [44, 45]. As these native molecular regulatory features may be

## Methods

### Tissue constructs

Primary articular chondrocytes were isolated from eight bovine carpometacarpal joints (3-6 months old) as previously described [46]. Cartilage tissues were digested in 1000 U/mL Type IV Collagenase (Worthington) overnight at 37 °C for 14 hours with orbital shaking. Isolated chondrocytes were seeded in 2% w/v agarose (type VII, Sigma) at 30 × 10^6^ cells/mL and cast into 2 mm-thick molds. Cylindrical tissue constructs were prepared via biopsy punch in a range of diameters (Ø2-Ø6 mm). Tissue constructs were maintained in chondrogenic media consisting of high glucose Dulbecco’s Modified Eagle’s Media limited or not present in engineered tissue systems, the controlled administration of physiologic doses of active TGF-β may be necessary to recapitulate activity levels experienced by cells *in vivo*, thus promoting similar beneficial developmental outcomes.

The adopted reduced-size construct model serves to establish the existence of TGF-β optimization profiles whereby the administration of physiologic doses can yield neocartilage with native-matched mechanical properties while mitigating hyaline cartilage developmental abnormalities (Fig. 8a). However, the repair of clinical chondral defects requires the fabrication of larger tissue specimens, which encounter significant limitations in the uptake of TGF-β supplemented in culture medium. Together, these characterizations point to the need for TGF-β scaffold delivery platforms capable of achieving sustained and uniform delivery of physiologic range TGF-β doses in order to yield the regeneration of uniform, large-size hyaline cartilage with optimized tissue quality. While the past decade has been marked by the emergence of a multitude of innovative TGF-β scaffold delivery platforms (e.g., microspheres, affinity domains, covalent conjugation [Fig. 8b]), they have predominantly been developed and tested for their utility in delivering highly supraphysiologic TGF-β doses to cells (Table 1). The results of this study advocate for tailoring scaffold delivery systems for achieving sustained delivery of physiologic TGF-β doses in order to improve cartilage regenerative outcomes.

**Fig. 8.**
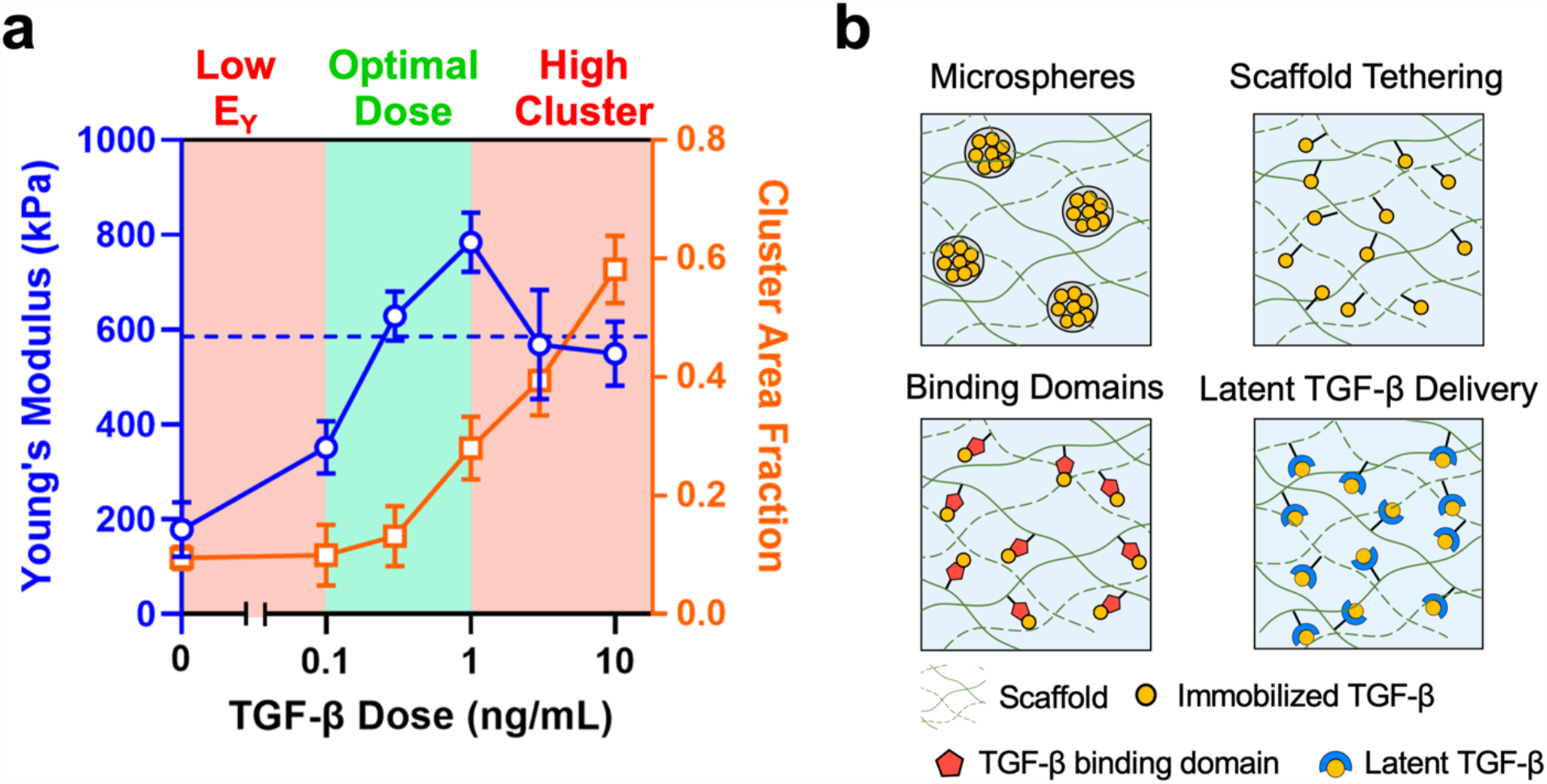
Future advances in TGF-β scaffold delivery platforms for optimization of cartilage regeneration. (**a**) Overlay of TGF-β dose-dependent curves for compressive Young’s modulus (E_Y_) (from Fig. 2) and cluster area fraction (CAF) (from Fig. 6). Optimized neocartilage consisting of native-matched mechanical properties while mitigating chondrocyte clusters can be achieved via TGF-β dose administration in physiologic range (0.1–1 ng/mL). Dashed line represents average E_Y_ of native bovine cartilage. (**b**) Optimized TGF-β dose delivery to construct-embedded cells can be achieved via emerging scaffold delivery platforms, including microspheres, scaffold tethering, affinity binding domains, and latent TGF-β delivery.

(DMEM, Gibco) supplemented with 1 mM sodium pyruvate, 50 mg/mL L-proline, 100 nM dexamethasone, 1% ITS+ premix (Corning), 1% PS/AM antibiotic-antimycotic, and 50 mg/mL ascorbate-2-phosphate (Sigma) at 37 °C with media refreshed three times per week over 56 days. For all studies, human recombinant active TGF-β3 (R&D Systems) was supplemented in culture media at specified concentrations for the initial 14 days of culture.

### TGF-β concentration gradient assessments

To measure TGF-β concentration gradients in constructs, Ø6×3 mm constructs (n=4) were exposed to 0.3 and 10 ng/mL TGF-β3 for 5 days. Constructs were subsequently axially sub-punched (Ø2 mm) and sectioned into 300-400 mm thick slices. The TGF-β3 content of each section was extracted via 4M Guanidine HCl (Sigma) at 4°C for 16 h and measured via ELISA (R&D Systems). The distribution of TGF-β in Ø2×2 mm and Ø5×2 mm constructs was predicted via FE model (FEBio) using the following reaction-diffusion equation (1) with previously measured governing parameters (Supplementary Table 1) [22]:

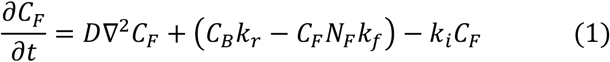

where *C*_*F*_ is the concentration of soluble TGF-β, *D* is TGF-β diffusivity, *C*_*B*_ is the concentration of bound TGF-β, *K*_*f*_ and *K*_*r*_ are the respective forward and backward reversible binding reaction rates, *N*_*F*_ is the concentration of the free binding sites, and *K*_*i*_ is the TGF-β internalize rate constant of chondrocytes.

### Mechanical property assessments

Construct compressive Young’s modulus (E_Y_) was measured using a customized testing device, as previously described [21]. Constructs (n=5 for day 56, n=3 for day 14, 28, 42) were subjected to 10% axial strain in unconfined compression and E_Y_ was computed via the equilibrium stress after 900 s of relaxation. For assessments of mechanical tissue heterogeneities, Ø3×2 mm and Ø5×2 mm constructs were exposed to 0.3 and 10 ng/mL TGF-β and were subjected to digital image correlation (DIC) strain analysis as previously described [47]. Briefly, diametrically-halved constructs (n=3) were stained over the exposed cross section with a speckled pattern of 1% hematoxylin (Sigma) via airbrush. The tissue cross section was imaged before and after the application of a 10% platen-to-platen compressive strain (Olympus CK40; 4× objective) and the spatial ε_yy_ strain map was assessed via Ncorr. Strain levels were averaged within six evenly divided axial sections through the tissue depth and normalized to the strain of media-exposed topmost tissue section.

### Biochemical composition assessments

Constructs were weighed and digested with 0.5 mg/mL Proteinase-K (MP Biomedicals) at 56 °C for 16 h. The DNA content was measured using PicoGreen dsDNA assay kit (Invitrogen), sGAG content was quantified via 1, 9-dimethylmethylene blue (DMMB) assay [48], and total collagen content was quantified via orthohydroxyproline (OHP) assay [49]. To assess ECM composition heterogeneities after 56 days of culture, Ø6×2 mm constructs exposed to 0, 0.3, and 10 ng/mL TGF-β were axially sub-punched (Ø2mm), sectioned (∼500 mm thick), and assayed for sGAG content. The swelling ratio (SR) was determined by the mass ratio of each construct between day 56 and day 0.

### Cell morphology analysis

Live constructs (n=6-10 for day 30, n=4-6 for day 56) were diametrically halved, sectioned (∼100 mm thick), stained with calcein AM (Invitrogen), and imaged over the exposed cross section via confocal microscopy (Olympus FV3000; UPLSAPO 20×). For cell cluster analysis, images were binarized using a superposition of Gaussian filtering [50] and active contouring [51] analysis to differentiate between cellular and ECM regions (Supplementary Information). To quantitatively distinguish between cells in a clustered versus isolated cell morphology, a dimensionless morphological factor (MF) was applied, as described:

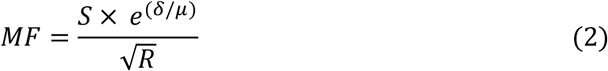

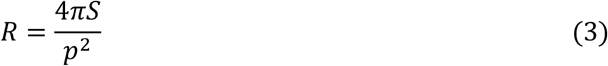

where *S* is the area of the cellular region, *δ* and *μ* is respective standard deviation and mean of cellular region fluorescence intensity, *R* and *p* is the roundness and perimeter of the cellular region, respectively. A frequency plot of the MFs of cellular regions from all analyzed images exhibited a bimodal distribution with a peak associated with isolated cell (MF=500) and clustered cell (MF=2000) associated peaks (Supplementary Fig. 5). A threshold value of MF=1500 was applied in order to distinguish between these regions. For each construct image, the cluster area fraction (CAF) was computed as the ratio of the area of clustered cell to total cellular area.

### Finite element analysis for cell strain

An FE model was implemented in MATLAB to predict minimum principal stress and strain of cells in native cartilage and TGF-β exposed constructs (0.3 or 10 ng/mL TGF-β) in response to applied physiologic loading. Models utilized binarized cell morphology images to define cellular and background ECM regions (Supplementary Fig. 4). Triangular mesh elements (maximum element size: 1 pixel^2^; minimum element size: 0.2 pixel^2^; number of elements: 825,000) and a linear elastic constitutive relation for cells and ECM was employed. Cellular region mechanical properties were prescribed from prior work (E_Y_ =1 kPa [52]; *v* =0.4 [53]). ECM region properties were prescribed from aforementioned bulk tissue stiffness measures (E_Y_; 0.3 ng/mL: 630 kPa; 10 ng/mL: 550 kPa; native cartilage: 590 kPa; Fig. 2) and prior work (*v*=0.2 [54]).

### Immunofluorescence

After 56 days of culture, constructs (n=5) were fixed in 10% formalin at 4 °C for 16 h, paraffin embedded, and sectioned. For Col-I, sections were predigested with 0.5 mg/mL hyaluronidase at 37 °C for 30 min and incubated in 0.5M acetic acid at 4 °C overnight [55]. For Col-II, sections were predigested with 0.1 mg/mL pronase (Sigma) at room temperature for 5 min. All sections were blocked with 10% goat serum for 30 min at room temperature. For Col-I staining, sections were incubated at 4 °C overnight with a 1:100 diluted mouse monoclonal anti-type I collagen antibody (C2456; Sigma). For Col-II staining, sections were incubated with a 1:128 diluted mouse monoclonal anti-type II collagen antibody (CIIC1; Developmental Studies Hybridoma Bank). All sections were incubated with goat anti-mouse Alexa Fluor 594 secondary antibody (1:500, Invitrogen) at room temperature for 1 hour and counterstained with DAPI (Mounting Medium With DAPI, Abcam). Images were acquired using a confocal microscope (Olympus FV3000; UPLSAPO 20×) and processed via ImageJ to quantify the fraction of Col-I positive cells.

### Statistical analyses

Two-way ANOVAs (α = 0.05) were performed to determine the effect of: 1) TGF-β dose and tissue depth (Fig. 1c), 2) TGF-β dose and construct size (Fig. 2a, Fig. 3a-b), and 3) TGF-β dose and culture duration on tissue properties (E_Y_, composition, or CAF) (Fig. 4a-b, Fig. 6b). One-way ANOVAs (α = 0.05) were performed to determine the effect of TGF-β dose on mechanical properties (Fig. 2b), biochemical contents (Fig. 3c-d), sGAG/collagen ratio (Fig. 3e), construct swelling (Fig. 3f) ratio, cell density (Fig. 4c), and positive Col-I staining cells (Fig. 5b). One-way ANOVAs (α = 0.05) were performed for the comparison of mechanical properties (Fig. 2a), sGAG content (Fig. 3a), and CAF (Fig. 6b) between tissue constructs and native cartilage tissue. Tukey’s HSD post-hoc tests were run to examine differences between experimental groups.

## Acknowledgments

Research reported in this publication was supported by the National Institute of Arthritis and Musculoskeletal and Skin Diseases under award number AR078299, the Boston University Dean’s Catalyst Award, the Boston University Material Science & Engineering Innovation Award, the Boston University Micro and Nano Imaging Facility and the National Institutes of Health under award Number S10OD024993, the Boston University Undergraduate Research Opportunities Program, and the Boston University Summer Term Alumni Research Scholars Program. The opinions, findings, and conclusions, or recommendations expressed are those of the authors and do not necessarily reflect the views of the National Institutes of Health. We thank Professor Paul E. Barbone for his insights and contributions to finite element modeling of tissue constructs in response to mechanical loading.

## Author contributions

T.W. and M.B.A. designed the study, performed the experiments, interpreted the data, and wrote the manuscript. S.D., Z.D., S.Y.K., and N.V. developed algorithms for cell morphology image analysis and cell strain finite element modeling. M.B.A. supervised the study, contributed to the scientific discussions, data interpretation, and to the manuscript.

## Data availability

The datasets generated during and/or analyzed during the current study are available from the corresponding author on reasonable request.

## Code availability

The steady state TGF-β distribution in tissue were simulated on FEBio (https://febio.org/). Images were analyzed on ImageJ (1.54f, https://imagej.nih.gov/ij/), Ncorr (v1.2, http://www.ncorr.com/), and MATLAB (R2022a). All codes are available upon reasonable request.

