## Supplementary Information for "Physiologic Doses of TGF-β Improve the Composition of Engineered Articular Cartilage"

#### Cell morphology analysis algorithm

We developed an algorithm for quantifying the degree of cell clustering in engineered articular cartilage tissues. The algorithm is composed of two parts: 1) image binarization to distinguish cellular regions from background, and 2) cell cluster determination (Supplementary Fig. 1). As the capacity to detect cell clusters is contingent upon the performance of the image binarization process, two independent image binarization techniques were applied to confocal images—Gaussian filtering and active contouring [1, 2]. The resulting binary matrices from each method were subsequently superimposed to yield a final matrix, which was used to spatially demarcate the area of cells with a clustered cell morphology versus those with an isolated morphology.

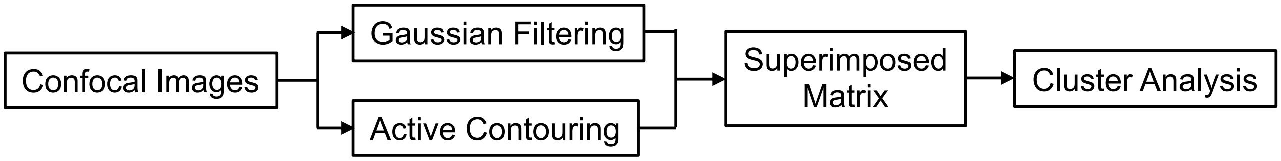


**Supplementary Figure 1.** Schematics of the cell morphology analysis algorithm.

#### 1. Image binarization

1.1 Gaussian filtering

A Gaussian filter is utilized on the confocal images in order to remove out-of-plane noise as well as homogenize, dilate, and merge cell regions in close proximity into a single transient region of foreground signal [3]. This allowed cell regions that were not necessarily spatially touching each other but still aggregating in a dense mass of cells to be blurred into a single cluster region [4]. Then a low-intensity threshold was applied to partition the foreground, fluorescent signals from those originating from out-of-plane cells. The threshold value was determined on the basis of the mean ($\underline{x}$) and standard deviation ($\sigma_x$) of the image intensity [5] by the following equation (1):

$$Low\ Intensity\ Threshold = \underline{x} + \alpha \cdot \sigma_x \qquad (1)$$

A coefficient value of α=1.0 was found to reduce the inclusion of out-of-plane fluorescence signals while retaining the dilated signals in the foreground of the Gaussian-blurred images without over-dilation (Supplementary Fig. 2).

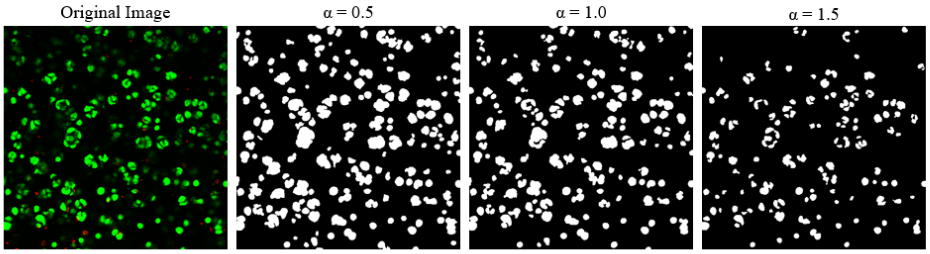


**Supplementary Figure 2.** The effect of coefficient in Gaussian filtering method on image binarization.

1.2 Active contouring

The second method employs the active contouring model developed by Chan, et al [6]. Briefly, the intensity gradient of the confocal images was first computed to initialize the contour curves outlining the cell regions. The curves were then evolved through the minimization of an energy-based segmentation model summarized in the equation (2) [7]:

$$F_1(C) + F_2(C) = \int_{inside(c)} |u_0 - c_1|^2 dx + \int_{outside(c)} |u_0 - c_2|^2 dx \quad (2)$$

The convergence criterion of active contouring is determined by the circumference of contour curve, where the contour iteration will stop when less than 1% difference in circumference is observed from the last iteration. The Chan-Vese active contours allowed us to detect cell regions without smoothing the initial image (Supplementary Fig. 3), thereby efficiently detecting while still preserving the boundary locations of the chondrocytes.

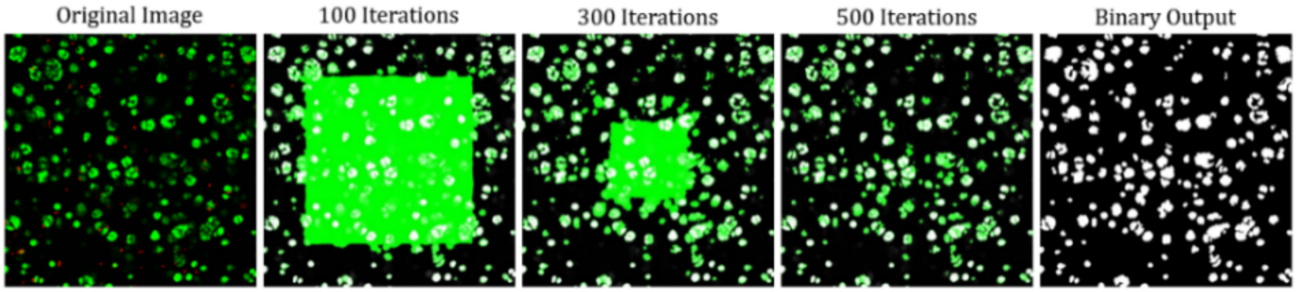


**Supplementary Figure 3.** Demonstration of the Chan-Vese active contouring for a confocal image.

### 2. Matrix superposition

Both Gaussian filtering and active contouring output a binary matrix that assigned a value of '1' to fluorescent signals of cell regions and '0' to the background. The two matrices were consequently superimposed into a single matrix, where a pixel value greater or equal to '1' signified that the pixel had been registered and validated as a foreground signal by at least one of the segmentation techniques. These pixels were preserved in the final superimposed matrix and subsequently used for morphology analysis (Supplementary Fig. 4).

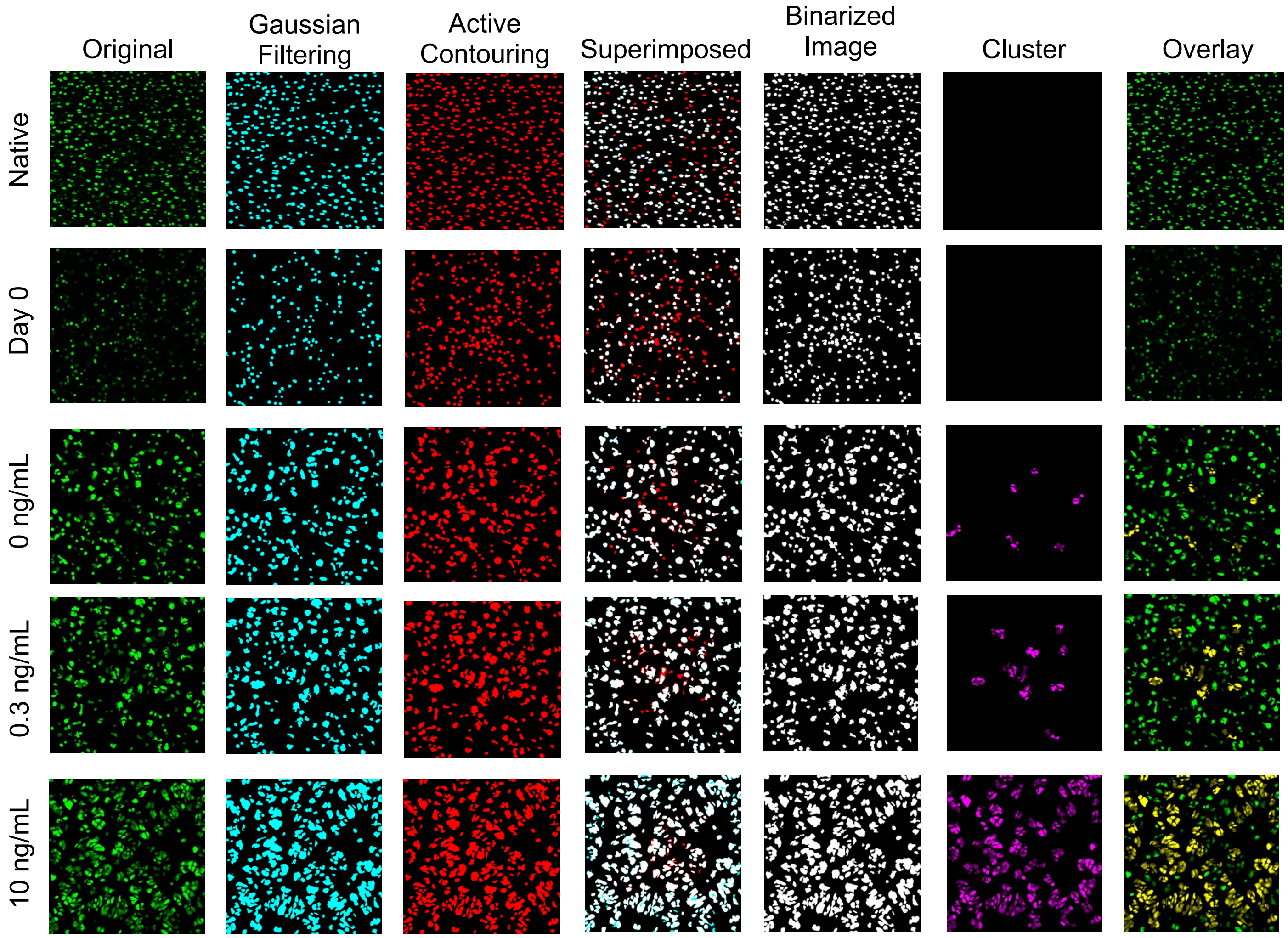


**Supplementary Figure 4.** Original image, binarization result of Gaussian filtering (cyan) and active contouring (red), superimposed output matrix of the two methods (white represents overlap of both methods), output binarized image (white), identified clustered cells (magenta), and overlay of clustered chondrocytes (yellow) and isolated chondrocytes (green) of native cartilage, day 0 construct, and day 56 constructs exposed to 0, 0.3, and 10 ng/mL TGF-β.

#### 3. Morphology analysis

Next, we defined a morphology factor (MF) to distinguish isolated and clustered cells [8, 9] described as the following equations (3-4):

$$MF = \frac{S \times e\hat{}(\delta/\mu)}{\sqrt{R}} \tag{3}$$

$$R = \frac{4\pi S}{p^2} \tag{4}$$

where $S$ is the area of the cellular region, $\delta$ and $\mu$ is respective standard deviation and mean of cellular region intensity, $R$ and $p$ is the roundness and perimeter of the cellular region, respectively. MF is a multifaceted term that accounts for many attributes of clusters that distinguish them from isolated cells. Clusters are formed with multiple cells and have larger areas than single cells, here, the diameter of a single chondrocyte was defined as 14 µm, determined by mean area of the chondrocytes in the confocal images of day 0 constructs, which is consistent to previous studies [10]. Further, isolated cells are more circular than cell clusters. Thus, roundness is one of the essential factors to define a cell cluster [8, 11]—low roundness values of clustered cells result in large MF. The coefficient of variation (CV), defined as the ratio between $\delta$ and $\mu$, indicates the distribution of the pixel intensities of each cellular region. Cell clusters formed by multiple cells tend to have higher CV due to the low-signal area in the gaps between cells. In contrast, isolated cells tend to have a low CV [12]. A frequency plot of the MFs of cellular regions from all analyzed images exhibited a bimodal distribution (Supplementary Fig. 5) with a peak associated with isolated cells (MF=500) and a peak associated with clustered cells (MF=2000). At the lower end of the spectrum, the high proportion of cell objects that have a MF lower than approximately 1500 are defined to be isolated chondrocytes. Hence, MF 1500 was determined as the cluster threshold—cell objects with MF higher than 1500 were defined as clusters (Supplementary Fig. 4). Based on this demarcation, a final cluster area fraction (CAF) was determined for each image, computed by the area of cell regions in a clustered morphology normalized to the total cell area.

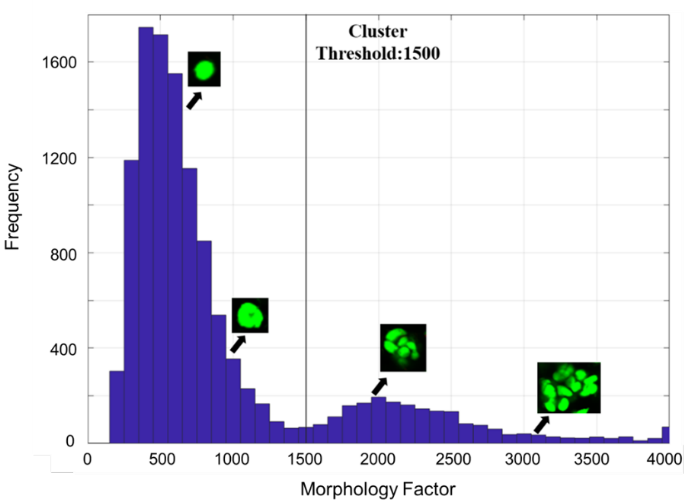


**Supplementary Figure 5.** Distribution of the morphology factor (MF) across all confocal images of constructs exposed to varying doses of TGF-β. MF 1500 is determined as the threshold for the identification of clustered cell regions.

**Supplementary Tables**

| Binding site density, $N_F$ [ng/mL] | Forward rate constant, $k_f$ [$\times10^{-6}$ s ng/mL$^{-1}$] | Reverse rate constant, $k_r$ [$\times10^{-5}$ s$^{-1}$] | Diffusivity, $D$ [$\mu m^2/s$] | Internalization rate, $k_i$ [$\times10^{-4}$ s$^{-1}$] |
|---|---|---|---|---|
| 200 | 2.7 | 2.5 | 23 | 4.5 |

**Supplementary Table 1.** Reversible binding, transport, and internalization parameters of active TGF-β3 in tissue constructs.
